# AHoJ: rapid, tailored search and retrieval of apo and holo protein structures for user-defined ligands

**DOI:** 10.1101/2022.09.05.496878

**Authors:** Christos P. Feidakis, Radoslav Krivak, David Hoksza, Marian Novotny

## Abstract

Understanding the mechanism of action of a protein or designing better ligands for it often requires access to a bound (holo) and an unbound (apo) state of the protein. Resources for the quick and easy retrieval of such conformations are severely limited.

Apo-Holo Juxtaposition (AHoJ) is a web application for retrieving apo-holo structure pairs for user-defined ligands. Given a query structure and one or more defined ligands, it retrieves all other structures of the same protein that feature the same binding sites(s), aligns them, and examines the superimposed binding sites to determine whether each structure is apo or holo, in reference to the query. The resulting superimposed datasets of apo-holo pairs can be visualized and downloaded for further analysis. AHoJ accepts multiple input queries, allowing the creation of customized apo-holo datasets. To demonstrate AHoJ’s functionality, we present a newly constructed dataset of apo-holo pairs featuring 13 ion ligands, by complimenting an existing database of biologically relevant holo interactions (BioLiP).

**Availability and Implementation:** Freely available for non-commercial use at http://apoholo.cz.

**Graphical abstract:** 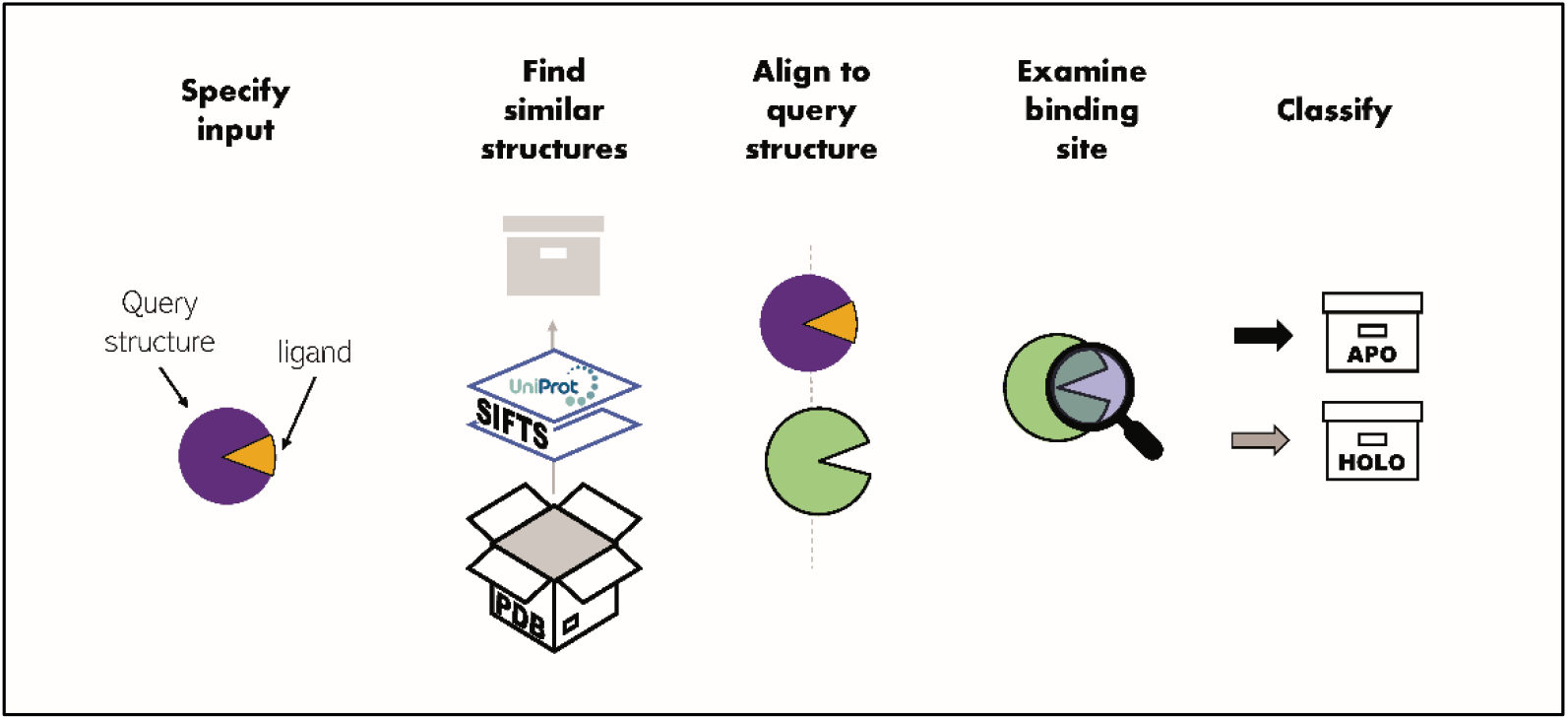

## Introduction and background

The study of protein–ligand interactions constitutes a prominent field in structural biology. Observing the effects of ligand binding (Brylinski and Skolnick, 2008), determining the temporal order of the conformational shifts that occur throughout the binding interaction (Morando et al., 2016), or exploring the promiscuity of a binding site (Ma et al., 2002), involve studying several protein– ligand interactions. Unveiling cryptic binding sites (Cimermancic et al., 2016), assessing the importance and consistency of water molecules (Wlodawer et al., 2018), or transcending the technical limitations of rigid body docking with ensemble docking methodologies (Amaro et al., 2018), also require access to several conformations. A common operational narrative among these topics includes the retrieval and comparison of multiple structures that involve a common point of interest, be that a ligand, binding site, or modified residue. Finding the relevant macromolecular structures in the PDB (Berman et al., 2000) is thus critical, but often cumbersome for structural biologists.

A number of datasets and tools have been built to address this need. ComSin (Lobanov et al., 2010) comprised a database of apo and holo protein pairs which exhibit significant shifts in their levels of intrinsic disorder upon complex formation. AH-DB (Chang et al., 2012) expanded this scope by including small ligands in its repertoire of apo-holo pairs. The BUDDY-system (Morita et al., 2011) provided a more flexible solution where the user could specify the ligand of interest, and the application would try to pair up the provided holo structure with an apo counterpart. At the time of writing, none of these servers are available. A recent work in preprint (APObind unpublished data) (Aggarwal et al., 2021) aims to complement an existing database of protein–ligand complexes, by pairing up the holo complexes with their apo counterparts. LigASite (Dessailly et al., 2008) is a more dated yet surviving resource that features pairs of apo and holo structures for 550 proteins. In both cases however, the ligand cannot be specified by the user.

The available resources appear to be restricted, and in some cases non-existent. The ability to define a ligand, and therefore a binding site, that will guide the search for apo and holo structures is missing altogether. This can be particularly useful as proteins often bind several ligands, and even within the same protein, different structures can bind different ligands in the same or in different binding sites. Therefore, finding pairs of apo and holo structures for a given target structure, requires specifying one or more ligands of interest. A methodology that defines the relevant ligands according to a fixed assumption (i.e., automatically), can restrict a user who wants to focus on a ligand that is deemed irrelevant, or narrow down the search to a single ligand when more bind the same structure. Ultimately, when an application forcefully decides upon the relevance of a ligand, it strips the user of this choice and it is also confronted with the non-trivial matter of biological relevance (Capitani et al., 2016).

Here, we present a web application that enables the user to conduct easy and fast parameterizable searches for apo and holo structure pairs against a target structure, by specifying one or more ligands of interest in this target structure, or letting the application detect the ligands instead. By probing the binding site of the user-defined ligand across other structures, it also allows exploring the binding preferences of the binding site and building a repertoire of ligands that natively bind the same site.

## Method outline

AHoJ starts the search by spatially marking the user-defined ligand(s) and identifying their binding residues with PyMOL. It then retrieves a list of candidate structure chains by i) detecting the UniProt accession number (AC) (UniProt: the universal protein knowledgebase, 2017) of each query chain and ii) reverse searching for all other chains that belong to the same UniProt AC. At the same time, it maps the binding residues of the query ligands onto the UniProt sequence by using the residue-level mappings from SIFTS (Dana et al., 2019), and cross-examines each candidate chain to determine how many of the mapped binding residues are present. If a minimum percentage of binding residues is detected, the chain is considered a successful candidate and it is aligned onto the query chain with TM-align (Zhang and Skolnick, 2005). The user can set the minimum percentage of mapped binding residues between query and candidate chain, as well as the minimum resolution, or the experimental method that was used to resolve the candidate structures (see S.I. for details on user parameters). The candidate’s area around the superimposed query ligand is examined for ligands, and the results are saved along with the aligned chains. This process is repeated for all candidate chains and each one is listed as holo or apo respective to the presence or absence of ligands in the defined binding site(s). In the presence of multiple defined binding sites in the query chain, if at least one of them is occupied by a ligand in the candidate chain, the chain is characterized as holo. The detected ligands along with metrics for similarity between candidate and query, presence of binding residues, and alignment scores, are reported for each apo and holo chain. The overall workflow is depicted in Fig. 1. Results are visualized in the browser and downloaded locally where they can be easily loaded into PyMOL through an included script.

**Fig. 1.**
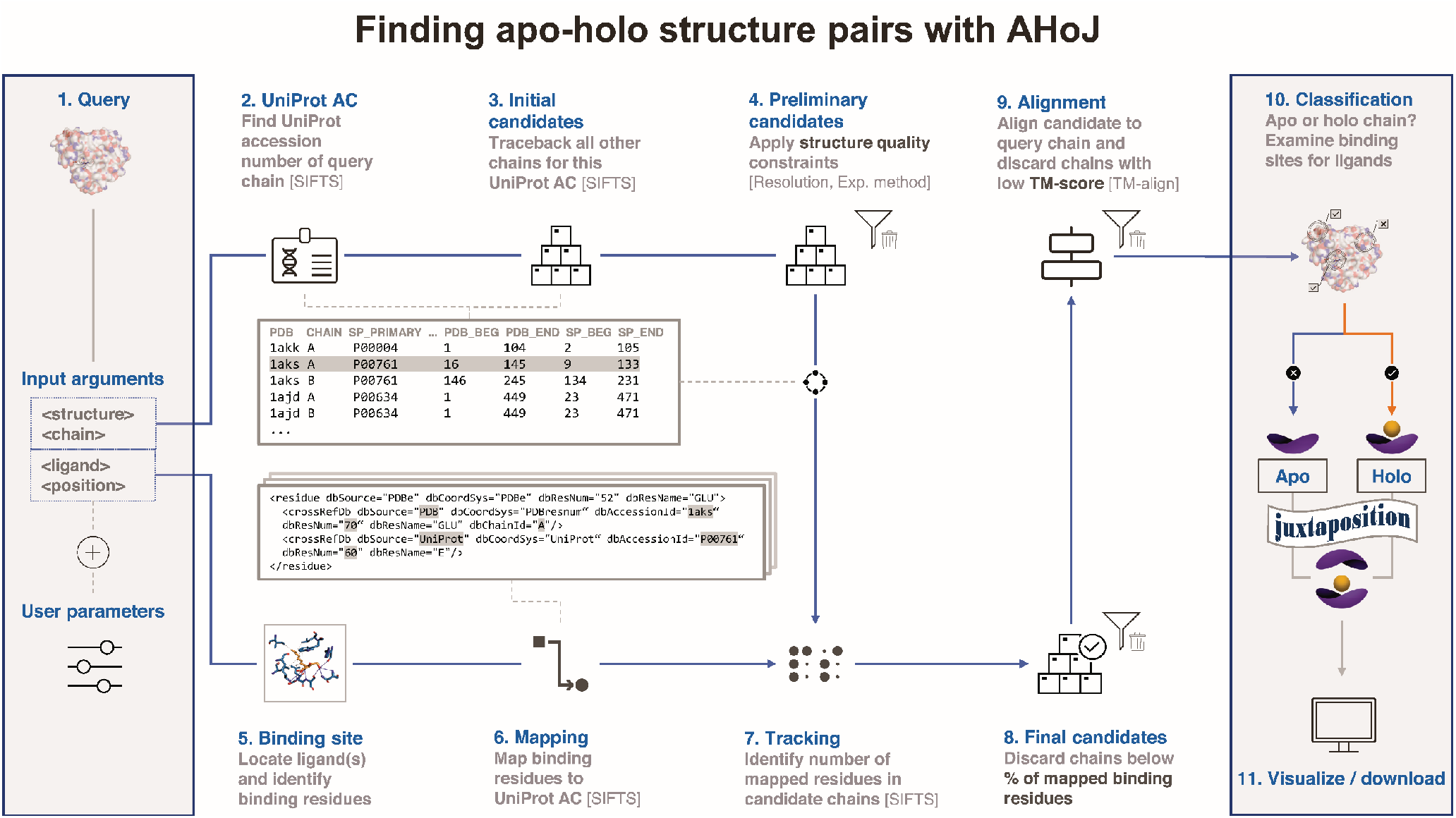
Flowchart depicting the workflow in AHoJ.

AHoJ’s main strength arises from allowing users to define the relevant ligand(s). Typically, ligands are confined to non-protein chemical moieties; however, in AHoJ the concept of ligand can be extended to include water molecules and modified or non-standard residues (e.g., phosphorylated residues or D-residues) as points of interest (see S.I. for details).

## Results and discussion

The idea to create AHoJ arose from our interest in ligand binding-site (LBS) prediction. We realized that testing structure-based LBS prediction applications exclusively on holo structures, can be unrealistic and produce results of questionable value. Nonetheless, such applications often use this approach or remain confined to limited apo datasets (Aggarwal et al., 2021). We therefore developed AHoJ to complement existing (holo) interactions with their apo counterparts and allow the creation of more realistic test and training sets for LBS prediction software.

To test AHoJ, we have chosen to complement a subset of the BioLiP database that features biologically relevant protein–ligand interaction information and is often used in drug design and machine learning applications (Yang et al., 2013). We have chosen 13 ion ligands, for which we extracted their interaction information from the non-redundant subset of BioLiP, totalling 38446 interactions across 11996 proteins (Table 1). These bound (holo) forms were used as input in AHoJ, which was able to retrieve unbound (apo) forms for 16987 of these holo interactions in 5080 proteins (approx. 44%). The raw results in AHoJ can feature several apo and holo chains, depending on how well each query protein is represented in the PDB. To create a practical dataset, we determined the top apo candidate for each initial (holo) interaction, by ranking the raw apo results of each search by the following sorting criteria (in order of impact): presence of the binding residues of the query (holo) structure in the apo structure (represented as a percentage), percentage of UniProt sequence overlap between the query and resulting apo chains, resolution and R-free of resulting structures (when available). The resulting apo-holo pairs are presented in the S.I.

**Table 1.** Names and number of interactions (sites) for 13 ion ligands (ligands are represented by their PDB naming convention). The bound (holo) forms were obtained from the BioLiP non-redundant set of biologically relevant ligands (Yang et al., 2013) and were used as input for AHoJ to search for their apo equivalent forms. Out of the 38446 interactions (holo), AHoJ was able to retrieve apo forms for 16987 of them (~44%). Site coverage refers to the percentage of holo interactions (sites) from the input that have at least one apo equivalent form. UniProt coverage refers to the percentage of proteins in the input that are represented in the retrieved apo-holo pairs.

| Ligand | Holo (input) |  |  |  | Apo (output) |  |  |  | Coverage (%) |  |
| --- | --- | --- | --- | --- | --- | --- | --- | --- | --- | --- |
|  | #UniProt | #structures | #chains | #sites | #UniProt | #structures | #chains | #sites | UniProt | sites |
| CA | 3184 | 5469 | 6975 | 10389 | 1433 | 2752 | 3415 | 4631 | 45 | 45 |
| CO3 | 111 | 180 | 244 | 290 | 64 | 115 | 154 | 183 | 58 | 63 |
| CU | 328 | 490 | 557 | 745 | 121 | 200 | 225 | 283 | 37 | 38 |
| FE | 520 | 803 | 1024 | 1206 | 128 | 237 | 328 | 375 | 25 | 31 |
| FE2 | 186 | 270 | 326 | 356 | 46 | 77 | 92 | 101 | 25 | 28 |
| K | 93 | 137 | 188 | 281 | 61 | 90 | 116 | 146 | 66 | 52 |
| MG | 3807 | 6316 | 8626 | 10480 | 2066 | 3476 | 4414 | 5395 | 54 | 51 |
| MN | 961 | 1539 | 2013 | 2589 | 495 | 839 | 1055 | 1343 | 52 | 52 |
| NA | 142 | 220 | 277 | 353 | 94 | 152 | 187 | 238 | 66 | 67 |
| NO2 | 36 | 83 | 103 | 135 | 17 | 47 | 56 | 81 | 47 | 60 |
| PO4 | 585 | 811 | 1142 | 1401 | 323 | 468 | 646 | 760 | 55 | 54 |
| SO4 | 504 | 630 | 937 | 1363 | 328 | 408 | 585 | 844 | 65 | 62 |
| ZN | 3982 | 5789 | 6674 | 8858 | 1140 | 1796 | 2098 | 2607 | 29 | 29 |
| 13 | 11996 | 21265 | 27820 | 38446 | 5080 | 7224 | 7800 | 16987 | 42 | 44 |

The same process can be employed for any subset of ligands or their entirety, giving rise to additional datasets of apo-holo pairs. Additionally, the raw results of apo and holo structures can be used to create ensembles, instead of utilizing a single structure. We hope that this work can support the transition to more complete and robust train and test datasets for machine-learning in the future.

## Supporting information

Apo-holo dataset

Raw results

## Acknowledgements

This research was supported by the Grant Agency of Charles University (project No. 1038120) and the ELIXIR CZ Research Infrastructure (ID LM2018131, MEYS CR).

