## Supplementary Information (S.I.) for "AHoJ: rapid, tailored search and retrieval of apo and holo protein structures for user-defined ligands"

#### Detailed methodology

AHoJ uses the UniProt accession number (AC) of a protein to build the original pool of candidates, and then leverages residue-level mappings between each PDB structure and its corresponding UniProt sequence (which are precalculated for every structure in SIFTS files), to measure their sequence overlap with the query sequence, and also to map the query binding site(s) across the candidate structures. These, along with an additional set of metrics, are used to measure the biological similarity between the candidate and the query structures. Some of these metrics are informative, meant to indicate the quality of the match (between candidate and query) and help users decide which resulting structures to use, and others are used as thresholds to filter out candidates that are deemed unsuitable for an apo or holo classification. The metrics are described herein according to their order of appearance in the application pipeline, along with the relevant user variables where applicable.

##### UniProt sequence overlap (mapping structures onto the UniProt sequence)

A key informative metric measures the percentage of overlap between the query structure chain and the prospective candidate chain. After all candidate structures for a given UniProt AC are retrieved (for a given query chain), each one is compared to the query sequence in terms of its overall coverage according to the start and end residue-level mappings that are available in the SIFTS files. This metric does not emerge from a pairwise alignment between candidate and query chain and does not refer to a sequence identity score; it is rather a comparison of the observed residues in each chain (i.e., present in the actual structures) that correspond to the same protein. The result is a percentage between 0 and 100, that reflects the percentage of amino acids in a given query structure chain, that are present in the given candidate structure chain. This metric is informative and it is not used as a cut-off threshold for filtering out candidate chains in a typical query where the query chain is holo, except for cases where the query chain is apo (and the default filtering by mapped binding residues cannot be applied). In such cases the user can specify a percentage as a minimum threshold (default is “0” (%)).
Note that this metric has directionality in the sense that the percentage of sequence overlap is computed from the perspective of the query chain and is therefore subject to length bias that may arise from comparing sequences of different sizes, much like in a typical sequence alignment. For

example, a long query chain would be less likely to have candidates with a high percentage of sequence overlap, while a shorter query chain would be more likely to have candidates with a high percentage of sequence overlap. This constitutes a basic incentive to avoid reliance on sequence overlap – and not use it as a candidate eligibility criterion when possible. AHoJ circumvents this in the case of holo query chains, by mapping the binding site(s) residues between query and candidate chains instead, whose presence or absence may be irrelevant to the overall sequence overlap (see "Mapping the binding site" for details).

#### Structure quality and experimental method

In AHoJ the user can specify a minimum resolution threshold which is applicable to structures that are resolved by scattering methods, in order to discard structures of unwanted resolution. Additionally, it is possible to exclude NMR structures or only consider X-ray crystallography structures. Note that in the latter case, structures resolved by hybrid methods including X-ray crystallography (e.g., electron paramagnetic resonance and neutron diffraction), will be excluded. These variables are used as thresholds for discarding structures. The R-free of the structures is reported in the results when available.

#### Mapping the binding site

When the query is a holo structure, AHoJ marks the position of the defined ligand(s) and identifies the binding residues (ligands need to be heteroatom entities according to the PDB file, see "Ligand detection" and "Notion of extended ligand" below for more information about ligand eligibility). It then looks for the presence of these binding residues in the candidate structures, to determine whether the candidate is suitable for an apo or holo assessment. This operation is performed by mapping the PDB numbering of the binding residues onto the UniProt sequence numbering and then cross-referencing these positions with the candidate structures, to determine if the residues are present or absent in the actual structure. This is performed by parsing the SIFTS files with the residue-level mappings of a given structure. The metric is used as a cut-off threshold to discard candidates that do not feature any of the binding residues of the query binding sites. A minimum cut-off of "1" (%) is set by default (user adjustable) and it is applied as the minimum percentage of binding residues that have to be present in the candidate chain out of the total binding residues in the query chain, for the chain to be classified as apo or holo. In the case of an apo query structure that does not bind any ligands and thus does not have a designated binding site, this metric is not applied. In such cases, the first metric (UniProt sequence overlap) is applied as a cut-off.

#### Alignment

The candidate chains that score above the previous threshold, are subsequently aligned to the query chain with TM-align (Zhang and Skolnick, 2005). This step also serves as a cut-off point, by specifying a minimum TM-score between the candidate and query chains (default = 0.5 (minimum TM-score), user adjustable). For each TM-align, two TM-scores are generated, each normalized by the length of the two aligned chains, which gives rise to inherent directionality -or length bias- depending on which chain is first and which is second. AHoJ captures both TM-scores generated in every alignment, and applies the minimum TM-score threshold (default = 0.5, user-adjustable) to the highest one, to avoid discarding candidate chains that score poorly on account of their low overall coverage against the query chain. The RMSD is also reported in the results as an informative metric.

#### Ligand detection

Successfully aligned candidates are assessed for ligands in the superimposed positions of the query ligands. Any heteroatom of the candidate structure that is positioned within a set radius (default = 4.5 Angstrom, user-adjustable) from the superposition of the specified (or auto-detected) query ligand atoms, is considered a ligand. By default, water molecules, modified residues and D-residues are ignored (user-adjustable). The user can also specify whether AHoJ should consider any detected ligand (within the above conditions) or restrict the search to the same ligand that was specified in the user query. This parameter is turned off by default, so that any detected ligand is considered in this step. If at least one ligand atom is detected within this scanning radius, the candidate structure is classified as holo, otherwise, as apo. In the presence of multiple defined binding sites in the query chain, if at least one of them is occupied by a ligand in the candidate chain, the chain is characterized as holo. The PDB names of the detected ligands are featured in the results for each candidate chain, and their positions (chain and PDB position index) are included in a separate CSV file with ligand information.

### The notion of extended ligand

AHoJ was designed around the premise that non-protein chemical moieties are the main point of interest in protein–ligand interaction research. Under this premise, it accepts any heteroatom as a ligand, that can be specified by its 1-3 PDB character code as a ligand name, and optionally also by its PDB position index in the structure.

Besides chemical compounds and ions, there is established evidence that water molecules hold a key role in understanding protein interactions (Schiebel et al., 2018). Furthermore, correctly assigning water molecules in the electron density maps of X-ray crystallographic structures, can be challenging, and has resulted in miss-annotations between water molecules and metal ions in deposited structures (Wlodawer et al., 2018). AHoJ allows users to define a water molecule as a ligand, and search for water molecules -or other ligands- in candidate structures in that particular superposition, in the same way that it would with a ligand, with the difference of changing internally the radius for scanning the candidate chain around the superposition of the query ligand from the default value of 4.5 to 2.5 Angstrom.

Another category of molecules that undoubtedly escape the definition of a ligand but are also important in understanding protein structure and function, are post translationally modified residues (e.g. phosphorylated residues). AHoJ allows users to specify such residue in a given structure, and search for apo and holo structures that lack or possess the specified modified residue in that particular superposition. Under the same principle, D-forms of amino acids can also be specified. Water molecules, modified residues and D-residues can be specified as input ligands through the user query or as candidate ligands (i.e., detectable entities affecting the apo or holo status of a candidate chain) through the respective parameters (*--water_as_ligand, --nonstd_rsds_as_lig, --d_aa_as_lig*).

### Usage

AHoJ works on the principle that users have a structure of interest and a point of interest on that structure (i.e. ligand, modified residue, water molecule) that they want to compare -in terms of the presence or absence of this point of interest- to the other structures of the same protein. The use-case can thus vary according to the user's input (type of point of interest) and the parameters, but the main objective is to perform comparisons for a given point of interest across different structures of the same protein. To accommodate this versatility in different types of points of interest, AHoJ offers a set of options through user-adjustable parameters and a text query format (single line input) that can accept 1 to 4 arguments.

#### Query format

The maximum arguments within the single line input are of this form:

<pdb_code> <chains> <ligand_name> <ligand_position>

- pdb_code: This is the 4-character code of a PDB protein structure (case-insensitive). This argument is obligatory and only 1 PDB code can be input per line. (e.g., "1a73" or "3fav" or "3FAV"). If it is the only argument (i.e., because the user does not know the ligand that binds to the structure), it will trigger automatic detection of ligands in the structure.

- chains: A single chain or multiple chains separated by commas (without whitespace), or "!" in the case of ligand-binding-only chains, or "*" in the case of all chains (i.e. "A" or "A,C,D" or "!" or "*"). This argument is case-sensitive and it is obligatory if the user intends to provide any argument after that (i.e. ligands or position).
- ligand_name: This argument is case-insensitive. A single ligand, multiple ligands separated by commas (without whitespace), or no ligands can be input per line (e.g., "HEM" or "hem" or "ATP" or "ZN" or "HEM,ATP,ZN") or "*" for the automatic detection of all ligands in the specified chain(s). Besides specifying the ligand directly by its name (and optionally, its position), the user can also specify a residue that binds the ligand (e.g., "HIS") and AHoJ will detect the ligand (as long as it is within 4.5 Angstroms of the residue). This approach however can lead to the selection of more than one ligands, if they are within this radius from the specified residue. This argument is non-obligatory, if omitted or specified as "*", AHoJ will automatically detect the ligands in the structure. If there are no ligands in the query structure, it will be characterised as apo and the search for candidates will continue. A water molecule can also be specified as a ligand (e.g., "HOH") but in such cases, its position must be specified as well. Note: when specifying the position argument, the user can only specify one ligand per query.
- ligand_position: This argument is an integer (e.g., "260" or "1"). It refers to the PDB index of the previously specified ligand, binding residue or water molecule. When this argument is specified, only one ligand or residue can be specified in the previous argument.

The primary mode of search in AHoJ, starts with a holo (bound) state. The most straightforward case is specifying a ligand as a point of interest. In such case, the ligand can be specified in the text query, by its 1-3 character PDB naming convention and also with its PDB index position in the amino acid sequence (this avoids considering all ligands of the same name that bind the same chain).

#### Examples

Example of a user query:
### consider ZN ligand in position 201 in chain A of PDB code 1a73 '1a73 A ZN 201'
The application will fetch the structure 1a73 and look for zinc+2 (ZN) ligand in chain A and position 201 of the sequence to validate the input. If ZN is found in chain A and position 201 of 1a73 (1a73A), it will retrieve all other known chains that belong to the same protein with 1a73A, align them with 1a73A and look for ZN (and also other ligands) at the superimposed binding site of ZN in 1a73A. If it finds protein chains with ZN, it will list them as HOLO, if the superimposed site

is empty of ligands, the chain will be listed as APO. If another ligand is detected on that site instead of ZN, the chain will be listed as APO or HOLO, depending on the value of *--lig_free_sites* parameter (if the user wants APO with no other ligands there, it will be listed as HOLO, and if the user allows other ligands in this binding site, it will be listed as APO).
Example of an alternative query that leads to the same result with the previous example:
### consider ligands near residue HIS134 in chain A of 1a73 (the detected ligand will be ZN 201 in chain A) '1a73 A HIS 134'

##### *More examples of user queries*

### consider ZN ligands in chains A and B of 1a73'1a73 A,B ZN' # consider ZN ligands in all chains of 1a73 '1a73 ALL ZN' or '1a73 * ZN' # find and consider all ligands in all chains of 1a73 '1a73' # find and consider all ligands in chain A of 1a73 '1a73 A' # consider ZN and MG ligands in chain A of 1a73 '1a73 A ZN,MG' # consider ZN ligands in all chains of 3fav '3fav ZN'

##### *Multiple queries*

Besides single queries, AHoJ also accepts multiple queries at once and processes them in batch mode. Queries are separated by line breaks, and one query is entered per line. The results for each query are saved in a separate folder and all of them are packed and downloaded in a single file. This can be useful for building datasets of apo and holo structures or simply processing multiple queries at once.
Example of a multiple query with comments for every single query (characters after “#” are ignored and can be used as comments):
1a73 A,B ZN # consider ZN ligands in chains A and B of 1a731a73 ALL ZN # consider ZN ligands in all chains of 1a731a73 # find and consider all ligands in all chains of 1a731a73 A # find and consider all ligands in chain A of 1a731a73 A ZN,MG # consider ZN and MG ligands in chain A of 1a733fav ALL ZN # consider ZN ligands in all chains of 3fav1DB1 # vitamin D3 study4est # porcine pancreatic elastase3CQV # reverb beta - all chains, all ligands3CQV A # reverb beta - chain A (in this case same effect)3CQV A HEM # reverb beta - ligand HEM (in this case same effect)

### Results

The results are visualized in the browser through Mol* and they can be downloaded as a zip file after a run has completed.

#### Files

In a successful run, AHoJ should generate the following files:

i) PDB structure files (cif.gz format) for the query structure (whole structure) and the successfully processed apo and holo candidate chains, aligned to the respective query chain(s).
Note: a given candidate chain could be a match for more than one query chains, and could thus appear more than once, in each case aligned to the respective query chain.
ii) 1 or 2 CSV files with the successfully processed candidate chains for apo and holo chains respectively [results_apo.csv, results_holo.csv]. These CSV files contain the following information for each found chain: **query_chain, apo_chain, Resolution, R-free, %UniProt_overlap, Mapped_bndg_rsds, %Mapped_bndg_rsds, RMSD, TM_score, iTM_score, ligands**
iii) 1 CSV file with the positions of the relevant ligands that were detected in both query and resulting candidate structures. This file is needed to load ligand selections when loading the results into the PyMOL with the included script.
Note: The ligands listed in the files refer to the ligands that were detected in the superimposed positions of the specified query ligands, thus they might not include ligands that bind elsewhere in the candidate chains. If the CSV file for apo chains includes ligands (which seems contradicting), it indicates that the user set the parameter *--lig_free_sites* to 0 (OFF), and thus any other ligands besides the query ligand were detected in the superimposed binding sites of candidate structures but ignored.
iii) 1 CSV file with information of the ligand positions for both query and candidate structures [ligands.csv]. This file is important for reference purposes and also if the user wants to reconstruct the PyMOL session with annotations locally.
iv) Console log file with information from the standard output [console.log]. This file can be used for reference and for better understanding the mechanism of action of AHoJ.
v) A PyMOL script file for loading the results into a PyMOL session [load_results_into_PyMOL.pml]. This is useful for viewing the results locally on the user's computer. The script has to be opened through PyMOL. The resulting session can then be saved by the user as a PyMOL session (.pse).

#### Visualization

i) The web application allows the visualization of the results in the browser with the molstar (Mol*) viewer. Web application: https://github.com/rdk/AHoJ-webapp
ii) The results can also be downloaded and visualized locally by loading the PyMOL script that is included in the results folder through PyMOL [load_results_into_PyMOL.pml]. The script has to be loaded from within the results folder. After downloading and unpacking the results into a folder, start a new PyMOL session and open the .pml file through it. A PyMOL installation is needed for this to work (Incentive or Open-Source)

### Parameters

#### Basic

**--res_threshold** : resolution threshold [default = 3.8]
Floating point number that represents angstroms and is applied as a cutoff point when assessing candidate structures that are resolved by scattering methods (X-ray crystallography, electron microscopy, neutron diffraction). It applies at the highest resolution value, when this is available in the PDB structure file. It can take any value, suggested min/max = 1.5/8. Condition is <=
**--include_nmr** : include NMR structures [default = 1]
0 or 1. When set to 1 (ON), NMR structures are considered as candidates. In the case of multiple states for a certain structure, the first one is considered.
**--xray_only** : x-ray structures only [default = 0]
0 or 1. When set to 1 (ON), only X-ray structures are considered. This overrides the NMR setting.
**--lig_free_sites** : ligand-free sites [default = 1]
0 or 1. When set to 1 (ON), it does not tolerate any ligands (in addition to the user-specified one(s)) in the superimposed binding sites of the candidate apo-proteins. When set to 0 (OFF), it tolerates ligands other than the user-specified one(s) in the same superimposed binding site(s). If the user wants to find apo structures that don't bind any ligands in the superimposed binding site(s) of the query ligand(s), they should set this value to 1 (default).

#### Advanced

**--bndgrsds_threshold** : binding residues threshold [default = 1.0, min/max = 1/100]
Floating point number that represents a percentage (%) and is applied as a minimum cut-off upon the percentage of the number of successfully mapped binding residues in the candidate chain out of the total number of binding residues in the query chain. The binding residues are mapped between query and candidate by converting PDB to UniProt numbering. "1%" translates to at least 1% percent of the query residues being present in the candidate structure, for the structure to be considered as apo or holo.
**--save_separate** : [default = 1]
0 or 1. When set to 1 (ON), the server will save all aligned chains that are in the opposite category from the starting query (apo/holo). In a regular search where the query is a holo-protein (searching for apo from holo), it will save any apo chains that it will find. In the opposite case when the query structure is apo, it would save all holo chains. If the user wishes to save both apo and holo chains, they can turn on the next parameter, "save_oppst".

**--save_oppst** : save opposite [default = 1]
0 or 1. When set to 1 (ON), the server will not only find, but also save chains that are in the same category with the starting query (apo/holo). In a regular search where the query is a holo-protein (searching for apo from holo), it will also save any holo chains that it will find. In the opposite case when the query structure is apo, it will also save the apo chains that it will find. This setting is dependent on the previous parameter "save_separate" which has to be ON for this parameter to work. This setting does not affect the search process of AHoJ which always includes both apo and holo chains.

**--overlap_threshold** : sequence overlap threshold [default = 0, min/max = 0/100]
Floating point number that represents a percentage (%) and is applied as a cutoff point when comparing the sequence overlap between the query and the candidate chain. It applies to the percentage of sequence overlap between query and candidate chains, and it is calculated from the query's perspective according to the UniProt residue numbering. If set to 100 (%), it means that the candidate chain has to completely cover the query chain. It can be longer than the query, but not shorter.

Note: "100" guarantees complete coverage, but it is the strictest setting. If the user wants a more lenient filtration, they can lower the value, or even set it to 0 and rely on the template-modeling score (TM-score) by using the default value (0.5) or setting their own TM-score cutoff with the "--min_tmscore" parameter.

**--lig_scan_radius** : ligand scanning radius [default = 4.0]
Floating point number that represents angstroms and is applied as a scanning radius when looking for ligands in the candidate structures. This scanning radius is applied on the positions of the atoms of the superimposed query ligands to the aligned candidate structure, to scan for ligands. The resulting scanning space is a "carved" surface that has the shape of the query ligand, extended outward by the given radius. If the candidate structure binds ligands outside of this superimposed area, they will be ignored, and the candidate will be characterised as an apo-protein.

**--min_tmscore** : minimum TM-score [default = 0.5, min/max = 0/1]
Floating point number that is applied as a minimum accepted template-modeling score between the query and the candidate chain. Value 1 indicates a perfect match, values higher than 0.5 generally assume the same fold in SCOP/CATH.

**--water_as_ligand** : [default = 0]
0 or 1. When set to 1 (ON), allows the detection of water molecules (i.e., 'HOH') as ligands in the superposition of the query ligand(s) in the candidate chains. If this setting is enabled and at least one water molecule is detected within the scanning radius, that would warrant a holo classification for

the candidate chain. When a water molecule is defined in the user query, this setting is automatically enabled.

**--nonstd_rsds_as_lig** : non-standard residues as ligands [default = 0]

0 or 1. When set to 1 (ON), allows the detection of non-standard -or modified- residues (e.g., 'TPO', 'SEP') as ligands in the superposition of the query ligand(s) in the candidate chains. If this setting is enabled and at least one modified residue is detected within the scanning radius, that would warrant a holo classification for the candidate chain. When a modified residue is defined in the user query, this setting is automatically enabled.

Note: The current list of non-standard residues includes the following residue names: 'SEP TPO PSU MSE MSO 1MA 2MG 5MC 5MU 7MG H2U M2G OMC OMG PSU YG PYG PYL SEC PHA'.

**--d_aa_as_lig** : D-amino acids as ligands [default = 0]

0 or 1. When set to 1 (ON), allows the detection of D-residues (e.g., 'DAL', 'DSN') as ligands in the superposition of the query ligand(s) in the candidate chains. If this setting is enabled and at least one D-residue is detected within the scanning radius, that would warrant a holo classification for the candidate chain. When a D-residue is defined in the user query, this setting is automatically enabled.

Note: The current list of D-residues includes the following residue names: 'DAL DAR DSG DAS DCY DGN DGL DHI DIL DLE DLY MED DPN DPR DSN DTH DTR DTY DVA'.
